# Transcriptomic and proteomic analysis show minimal role for mast cell ST2 in primary *Heligmosomoides polygyrus bakeri* infection

**DOI:** 10.1101/2025.11.03.686316

**Authors:** Nicole W P Ong, Suzanne H Hodge, Christa P Baker, Danielle J Smyth, Tania Frangova, Henry J McSorley

## Abstract

The IL-33/ST2 pathway is important as part of the type 2 immune response against helminth infections. Mast cells express the highest levels of the IL-33 receptor subunit ST2 of any immune cell, and mast cells can mediate type 2 immune inflammation, however the role of IL-33-driven mast cell responses in helminth infection is poorly understood. We sought to determine the role of mast cell ST2 expression during *Heligmosomoides polygyrus bakeri* (*Hpb)* infection by generating mast cell conditional ST2 knockout (MCPT5^C^re x ST2^f/f^, cKO) mice. These mice have normal frequencies of mast cells at steady state, but show specific and strong (albeit incomplete) knockdown of ST2 expression on mast cells. On *Hpb* infection, faecal egg and adult worm burden were similar between cKO and littermate controls, as were mast cell degranulation markers, serum IgE and goblet cell hyperplasia. Therefore, we conclude that mast cell ST2 does not play a dominant role in *Hpb* infection. To further investigate the immune response to infection in cKO and littermate controls, transcriptomic and proteomic changes were assessed in duodenal tissues in infected versus naïve mice in cKO and control mice. Minimal transcriptomic and proteomic changes were seen between genotypes, whereas substantial changes were seen between naïve and infected mice, regardless of genotype. *Hpb* infection induced local increases at the transcript and protein level for mast cell proteases (MCPT1 and MCPT2), resistin-like molecules (RELMα and RELMβ), and markers such as the phospholipase PLA2G4C and the pore-forming protein gasdermin C. Bulk proteomic analysis was also searched against the *Hpb* genome to identify *Hpb* proteins present in the duodenal tissues. A list of 60 *Hpb* proteins of interest were identified in infected duodenal samples, of which 18 contain a signal peptide and are present in the excretory/secretory products of *Hpb* (HES) (likely secretory products including immunomodulatory proteins); 28 proteins are present in HES but do not contain a signal peptide (likely excretory products); and 14 proteins are not present in HES (likely proteins present in the remnants of *Hpb* within the duodenum). This work thus provides datasets for changes in the mouse intestine due to *Hpb* infection, at both the transcript and protein level, as well as a dataset of *Hpb* proteins detectable in the mouse duodenum at day 14 of infection.

## Introduction

Soil transmitted helminths (STH) infect hundreds of millions of people worldwide, causing significant morbidity including anaemia, impaired growth, and gastrointestinal manifestations of ill-health (WHO, 2023). They are often associated with a type 2 immune response, characterised by the release of epithelial cytokines such as IL-25 and IL-33, activation of innate and adaptive lymphocytes such as type 2 innate lymphoid cells and CD4+ Th2 cells; the release of type 2 effector cytokines including IL-4, IL-5 and IL-13; the induction of IgE antibody responses; and the recruitment and activation of type 2 effector cells such as eosinophils and mast cells (Reynolds et al., 2012; Zaiss et al., 2024). These type 2 immune responses also result in changes to the architecture of the intestine, through acting on stromal cells to induce proliferation of secretory cells such as goblet cells, increased epithelial turnover, and smooth muscle contraction, leading to the so-called “weep and sweep” response which can eject parasites from the intestinal lumen (Grencis, 2015).

Parasitic helminths have developed a range of effector molecules to suppress the development of the types of immune responses that eject them. In the case of the model intestinal nematode *Heligmosomoides polygyrus bakeri* (*Hpb*), these include the secretion of various immunomodulatory protein families including the HpARIs (which bind IL-33) (Osbourn et al., 2017), the HpBARIs (which bind the IL-33 receptor, ST2) (Vacca et al., 2020), the TGMs (which bind the TGF-β receptor) (Johnston et al., 2017) and HpBoRB (which binds to the anthelmintic effector protein RELMβ) (Shek et al., 2025). As both the HpARIs and the HpBARIs block the IL-33 pathway, it follows that this cytokine pathway is particularly important to resistance to intestinal nematodes. Indeed, total IL-33- or ST2-deficient mice show increased susceptibility to a wide range of intestinal helminths, including *Hpb* (Coakley et al., 2017; McSorley and Smyth, 2021). However, the proximal responders to IL-33 release in the intestine, and the cellular target of HpARI- and HpBARI-mediated suppression, is not well understood.

There are multiple cell types which express ST2, the cognate subunit of the IL-33 receptor, including ILC2s, Th2 cells and eosinophils, however the immune cell type with the highest expression of ST2 is the mast cell (Immunological Genome, 2020; Brenes et al., 2023). When mast cells are stimulated with IL-33, they secrete cytokines such as IL-6 and IL-13, and IL-33 stimulation can synergise with FcERI-mediated degranulation to further increase cytokine release (Silver et al., 2010). Despite their association with IL-33 responsiveness and IgE-FcERI-mediated degranulation responses, the role of mast cells in helminth infection is under-researched. Mast cells have been implicated in the production of IL-33, either cell-intrinsically (Shimokawa et al., 2017), or via induction of the release of epithelial cell-derived IL-33 (Yang et al., 2024). However, their role in responding to the cytokine is unclear.

We hypothesised that mast cell responses to IL-33 could be important in the type 2 immune response to the parasite. To this end, we used MCPT5^Cre^ x ST2^f/f^ (cKO) mice to determine the role of IL-33 signalling in mast cells during *Hpb* infection. We assessed immune responses in the draining lymph nodes, and transcriptomic and proteomic studies of the intestine to assess responses to the parasite. Finally, we used proteomics to identify parasite proteins which are retained in the intestinal tissue once the parasites have left the wall of the gut.

## Methods & Materials

### Mice & Ethics Statement

Wild-type C57BL/6J were purchased from Charles River, UK. *Mcpt5-Cre* mice (C57BL/6J background) (known henceforth as “MCPT5^Cre^”) (Scholten et al., 2008) were provided by Prof. Dr. med. Axel Roers (Heidelberg University, Germany). C57BL/6N-A^tm1Brd^ *Il1rl1*^tm1c(KOMP)Wtsi^/RleeMmucd mice, RRID:MMRRC_048182-UCD (known henceforth as “ST2^f/f^”), were obtained from the Mutant Mouse Resource and Research Center (MMRRC) at University of California at Davis, an NIH-funded strain repository, and was donated to the MMRRC by The KOMP Repository, University of California, Davis; Originating from Dr Richard Lee, Brigham and Women’s Hospital (Skarnes et al., 2011). MCPT5^Cre^mice were crossed with ST2^f/f^ mice, then backcrossed to give required genotypes. MCPT5^C^re x ST2^f/f^ mice were used as conditional knockouts (cKO), while MCPT5^WT^ x ST2^f/f^ littermates were used as controls. Mice were housed at 19-21°C, 45-65% humidity, under a 12/12-hour light dark cycle and allowed *ad libitum* access of tap water and commercial RM3 pellet food. Mouse accommodation and procedures were performed under UK Home Office license PP9520011 with institutional oversight performed by qualified veterinarians.

### *Hpb* Maintenance, Infection and Parasitological Readouts

Life cycle of *Hpb* was maintained as described previously (Johnston et al., 2015). Mice were infected with *Hpb* by oral gavage of 200 L3 *Hpb* in water. Adult *Hpb* worm burden was quantified by dissecting whole small intestine longitudinally and manually picking out *Hpb* worms. *Hpb* egg burden was quantified by collecting and weighing faecal pellets, which were then soaked in water and resuspended in a saturated salt solution to create a faecal slurry. Faecal slurry was then deposited on a McMaster slide for *Hpb* egg counts.

### Serum Collection, Peritoneal Lavage (PL), Tissue Dissection and Preparation of Single Cell Suspensions

Mice were culled by overdose of medetomidine and ketamine, and exsanguinated by severing of a major artery to collect blood for serum analysis. The peritoneal cavity was lavaged with two washes of 5 mL of ice-cold PBS to collect peritoneal lavage (PL) cells. Dissected mesenteric lymph nodes (mLNs) were macerated through a 70 μm cell strainer (BD Biosciences) in RPMI 1640 medium to obtain a single cell suspension. Red blood cells in PL were lysed by incubation with ACK Lysing Buffer (Gibco™) at room temperature before quenching and centrifugation at 400g, 5 mins. Supernatant was discarded and PL or mLN cells were resuspended in PBS or FACS buffer (PBS supplemented with 0.5% BSA and 0.05% sodium azide) respectively for cell counts with a haemocytometer.

### Flow Cytometry

For surface staining, cells were incubated with Zombie UV™ Fixable Viability dye (Biolegend) for 20 mins in the dark, at room temperature, then washed and incubated with anti-CD16/32 (Biolegend) in FACS buffer (PBS supplemented with 0.5% BSA and 0.05% sodium azide) for 10 mins in the dark, at 4°C. Cells were washed and incubated FACS buffer containing appropriate antibodies for 20 mins in the dark, at 4°C. Antibodies used for surface staining were a lineage cocktail either on FITC (CD3-FITC (clone 145-2C11), CD5-FITC (clone 53–7.3), CD11b-FITC (clone M1/70), CD19-FITC (clone 6D5), CD49b-FITC (clone DX5, Invitrogen), GR-1-FITC (clone RB6-8C5)) or Pacific Blue (CD3-Pacific Blue (clone 17A2), CD5-Pacific Blue (clone 53–7.3), CD11b-Pacific Blue (clone M1/70), CD11c-Pacific Blue (clone N418), B220-Pacific Blue (clone RA3-6B2), NK1.1-Pacific Blue (clone PK136); CD45-AF700 (clone 30-F11) or CD45-APCCy7 (clone I3/2.3), ckit-Pacific Blue (clone 2B8), FcεRI-PECy7 (clone MAR-1), IgE-PE (clone RME-1), CD127- FITC (clone A7R34), CD90.2-AF700 (clone 30-H12), CD4-Brilliant Violet 711 (clone GK1.5), ST2-APC (clone RMST2-2, Invitrogen) or ST2-biotin (clone DIH9) and Streptavidin-APC (all from Biolegend except where indicated). Cells were washed and resuspended in FACS buffer prior to acquisition. Acquisition was performed on the LSRFortessa™ (BD Biosciences) and data was analysed on FlowJo software v10 (BD Life Sciences, FlowJo LLC).

For intracellular staining, eBioscience™ Foxp3 / Transcription Factor Staining Buffer Set (Invitrogen) was used as per manufacturer’s instructions. Briefly, after surface staining, cells were incubated in Foxp3 Fixation/Permeabilisation solution overnight at 4°C in the dark. Cells were incubated with 1x permeabilisation buffer containing antibodies against GATA3-PE (clone TWAJ, Invitrogen) and Foxp3-PECy7 (clone FJK-16s, Invitrogen), for 30 mins in the dark, at 4°C prior to washing and acquisition.

### ELISA

For MCPT-1 (Invitrogen), MCPT-4 (Aviva Systems Biology) and IgE (Biolegend), ELISA kits were used as per manufacturer’s instructions. OD450 and OD570 readings were taken with a CLARIOstar Microplate Reader (BMG Labtech).

### Histology and Imaging

Small intestines were prepared for histology as described previously (Smyth et al., 2025). Briefly, small intestines were dissected out and contents flushed with PBS, fixed in 4% neutral buffered formalin and rolled into Swiss rolls. After tissue processing and embedding in paraffin, 5 μm sections were cut and stained with Periodic Acid-Schiff (PAS). Samples were visualised under a Leica ICC50 HD microscope, with images were taken with Leica LAS EZ software and analysed on Fiji software. Cell counts were performed manually and intestinal area was quantified by area calculation of manually indicated regions on Fiji 1.53q (open-source (Schindelin et al., 2012)).

### Bulk-RNA Sequencing

The first 0.5 cm of small intestine from the duodenum end was dissected and stored in RNAlater solution (Thermo Fisher) overnight at 4°C, then stored at -70°C until processing and data analysis by Azenta Life Sciences whereby tissues were used for bulk RNA sequencing with PolyA section, 30 million read pairs per sample. Read counts were normalised across samples to determine differentially expressed genes. DESeq2 was used to compare gene expression across treatment groups with the Wald test and Benjamini-Hochberg procedure used to calculate q-values. Genes with a q-value <0.05 and absolute log2 fold change >1 were determined as significantly differentially expressed genes.

### Sample Preparation for Mass Spectrometry

Subsequent 0.5 cm of small intestine from the duodenum end was dissected, flash frozen in dry ice and stored at -70°C until processing for mass spectrometry. Sample preparation protocol was adapted from published literature (Baker et al., 2022). Briefly, tissue was digested in lysis buffer (5% SDS, 50mM TEAB in water) and sonicated at maximum amplitude for 15 cycles of 30 seconds on, 30 seconds off. Tubes were briefly centrifuged to collect liquid to bottom of tubes, and benzonase (Merck) added. Protein yield was determined using BCA method, then TCEP (final concentration 10 mM) added and incubated at 55°C for 5 min. 0.5M iodoacetamide (Sigma) was added to samples and incubated at room temperature for 1 hour in the dark. Lysates were acidified and loaded onto S-Trap™ mini spin columns (ProtiFi) for trypsin (Promega) digestion and peptide dehydration as described previously (Baker et al., 2022). A sample of excretory/secretory products of *Hpb* (HES – prepared as previously described (Johnston et al., 2015)) was also prepared and run for proteomic analysis and comparison.

### Liquid Chromatography-Mass Spectrometry (LC-MS)

Mass spectrometry was conducted by University of Dundee FingerPrints Proteomics Facility. Sample volumes equivalent to 1.25 µg of peptides were injected onto an UltiMate™ 3000 RSLCnano system (Thermo Scientific) and electrosprayed into an Orbitrap Exploris 480 Mass Spectrometer (Thermo Fisher). The following LC buffers were used: Buffer A - 0.1% formic acid (Fisher Scientific) in MilliQ water (v/v)); Buffer B - 80% acetonitrile (VWR) and 0.1% formic acid in MilliQ water (v/v). Samples were loaded at 10 µl/minute onto a trap column (100 µm x 2 cm, PepMap nanoViper C18 column, 5 µm, 100 Å, Thermo Scientific) equilibrated with 0.1% trifluoroacetic acid (TFA, Thermo Scientific). The trap column was washed for 3 minutes at the same flow rate with 0.1% TFA then switched in-line with a resolving C18 column (75 μm × 50 cm, PepMap RSLC C18 column, 2 μm, 100 Å, Thermo Scientific). Peptides were eluted from the column at a constant flow rate of 300 nl/min with a linear gradient from 3% buffer B to 6% buffer B in 5 minutes, then from 6% buffer B to 35% buffer B in 115 minutes, and finally from 35% buffer B to 80% buffer B within 7 minutes. The column was then washed with 80% buffer B for 4 minutes. Two blanks were run between each sample to reduce carry-over. The column was kept at a temperature of 50 °C.

The data was acquired using an easy spray source operated in positive mode with spray voltage at 2.25 kV, and the ion transfer tube temperature at 250 °C. The MS was operated in DIA mode. A scan cycle comprised a full MS scan (m/z range from 350-1650), with RF lens at 40%, AGC target set to custom, normalised AGC target at 300%, maximum injection time mode set to custom, maximum injection time at 20 ms, microscan set to 1 and source fragmentation disabled. MS survey scan was followed by MS/MS DIA scan events using the following parameters: multiplex ions set to false, collision energy type set to normalized, HCD collision energies set to 25.5, 27 and 30%, orbitrap resolution 30000, first mass 200, RF lens 40%, AGC target set to custom, normalized AGC target 3000%, microscan set to 1 and maximum injection time 55 ms. Loop control N, N (number of spectra set to 23). Data for both MS scan and MS/MS DIA scan events were acquired in profile mode.

### Proteomic Analysis

Raw data files were ran on Spectronaut 19 against *Mus musculus* (Swissprot Trembl November 2023), *Hpb* (transcriptome described previously (Hewitson et al., 2011)) and common contaminants fasta file using stringent identification settings (Baker et al., 2024). Intensity-based absolute quantification (iBAQ) was exported from Spectronaut and used for quantitative analysis. *M. musculus* and *Hpb* proteomes were separated, and differential comparisons were conducted using limma function outputs including log2 fold changes, p-values, and q-values (Ritchie et al., 2015). *Hpb* proteins were analysed for the presence of a signal peptide using SignalP - 6.0 (Teufel et al., 2022).

### Statistical Analysis

Statistical analysis was performed on Graphpad Prism 10 (Graphpad). For comparisons across two groups, unpaired t-test was used. For comparisons across more than two groups, one-way ANOVA or two-way ANOVA and multiple comparisons was used as stated in figure captions. Reports of statistical values are as follows: ns (p > 0.05), * (p ≤ 0.05), ** (p ≤ 0.01), *** (p ≤ 0.001), **** (p ≤ 0.0001).

### Data Availability

Bulk-RNASeq data will be uploaded to the Gene Expression Omnibus, while proteomics raw files, Spectronaut reports, FASTA files and experiment templates will be uploaded to PRIDE (Perez-Riverol et al., 2022) on acceptance of the manuscript.

## Results

### MCPT5^C^re x ST2^f/f^ mice show no change in susceptibility to *Hpb* infection

To investigate the role of ST2 expression on mast cells, we generated MCPT5^C^re x ST2^f/f^ (cKO) mice to conditionally delete ST2 in MCPT5-positive mast cells. On *Hpb* infection, we found that these cKO mice showed no change in egg or parasite burden over 28 days of infection, when compared to Cre-negative littermate controls (**Figure 1A-B**). Immunological responses to *Hpb* were assessed at day 14 of infection (at the peak of the type 2 immune response) showing similar induction of serum IgE and mast cell proteases MCPT-1 and MCPT-4 in cKO and control mice (**Figure 1C-E**). This suggests that there was no difference in IgE-mediated mast cell degranulation in cKO mice compared to controls.

**Figure 1:**
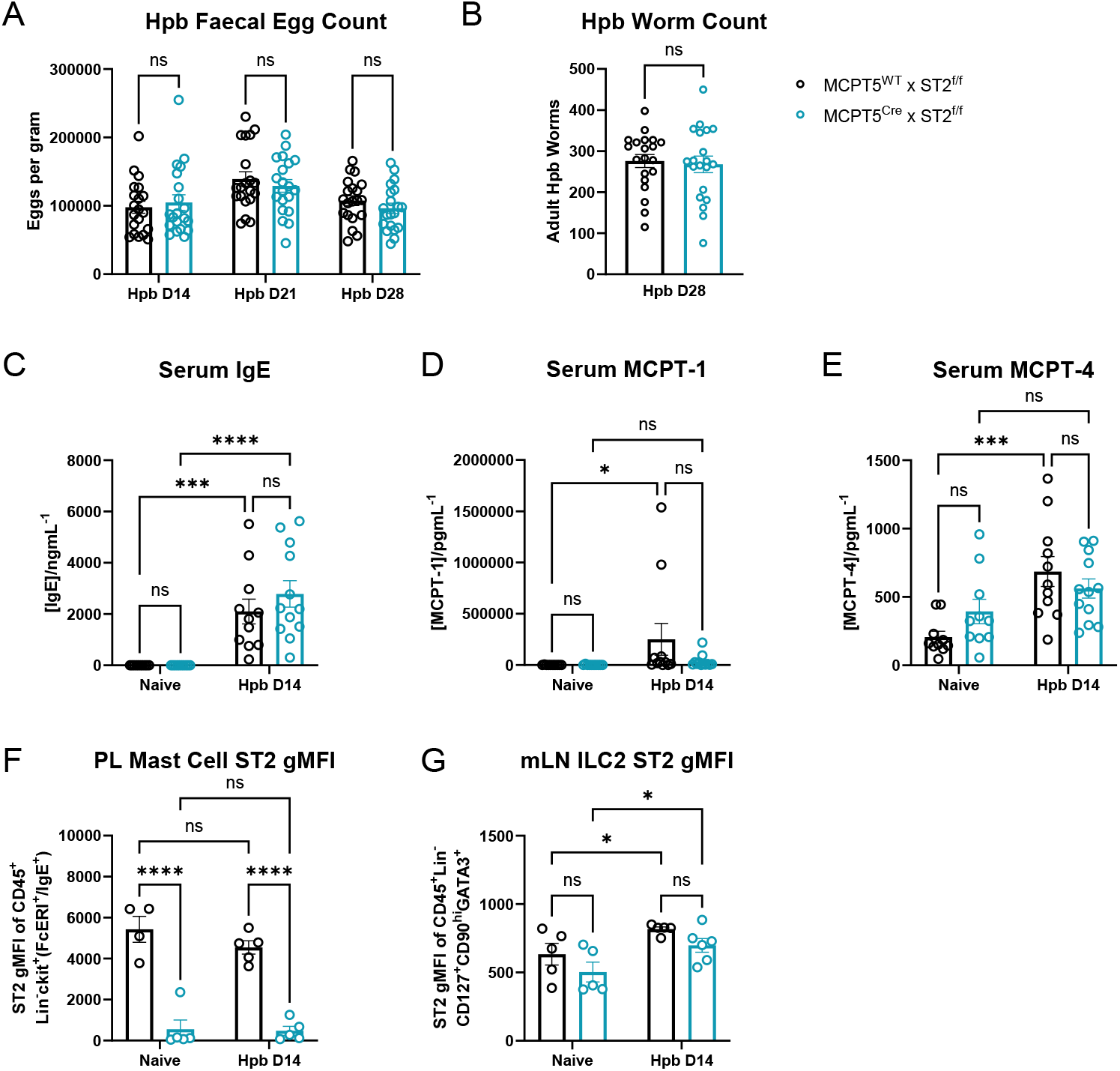
Mast cell ST2 plays no role in susceptibility and serum responses to *Hpb* infection. MCPT5^Cre^ x ST2^f/f^ and MCPT5^WT^ x ST2^f/f^ mice were infected with 200 L3 *Hpb* larvae by oral gavage. At days 14, 21 and 28 of infection *Hpb* eggs per gram of faeces (**A**) and adult worm burdens in the small intestine at day 28 of infection (**B**) were assessed. Data in (**A-B**) is pooled across 2 independent experiments, each with n=10/group and was analysed by 2- way ANOVA with Tukey’s multiple comparisons test. Separate cohorts of mice were culled at day 14 of infection (versus naïve controls) and serum levels of total IgE (**C**), mast cell protease-1 (MCPT-1) (**D**) and MCPT-4 (**E**) were measured. Data in (**C-E**) is pooled across 2 independent experiments, each with n=4-5/group and was analysed by Welch’s t-test. Peritoneal lavage (PL) and mesenteric lymph node (mLN) cells were also prepared for flow cytometry to assess mast cell (**F**) and type 2 innate lymphoid cell (ILC2) (**G**) ST2 expression (geometric mean fluorescence intensity (gMFI)). Data in (**F**-**G**) is representative of 2 experiments, each with n=4-5/group and was analysed by 2-way ANOVA and Fisher’s LSD test. Mice were 8-20 weeks old, with both males and females used. Error bars show mean ± SEM.

In the course of this work, it became clear that staining for FcERI (used to gate on mast cells) varied during *Hpb* infection, and that over a timecourse of *Hpb* infection of wildtype mice, the FcERI signal by flow cytometry negatively correlated with total IgE levels (**Supplementary Figure 1A-D**). This effect was attributed to an artefact of the inability of the MAR-1 clone of the anti-FcεRI antibody to bind IgE-bound FcεRI (Khodoun et al., 2013; Worrall et al., 2022). Therefore, FcεRI is an unreliable marker of mast cells during infection, and subsequently mast cells were gated as CD45^+^Lin^-^ckit^+^(FcεRI^+^/IgE^+^), thereby including cells that were singly positive for either FcεRI or IgE to account for this artefact (**Supplementary Figure 1E**).

Taking this into consideration, ST2 expression on mast cells at day 14 of *Hpb* infection was similar to that seen in naïve mice, with a robust deletion of ST2 expression in mast cells from cKO mice (**Figure 1F**), while mesenteric lymph node (mLN) ILC2s retained normal levels of ST2 expression (**Figure 1G**). Therefore, these cKO mice showed a specific knockdown of ST2 on MCPT5-positive mast cells, and not other immune cells, as expected.

Type 2 immune responses in the mLN were also measured at day 14 of infection. Th2 and ILC2 responses also appeared unaffected, with similar robust induction of GATA3^+^Foxp3^−^ Th2 frequencies of CD4^+^ T cells, and total Th2 numbers on infection, in both cKO and control mice (**Figure 2A-B**). *Hpb* infection increased GATA3^+^Lineage^−^ ILC2 numbers, but not frequency, and this response was again unaffected in cKO versus littermate controls (**Figure 2C-D**). Finally, goblet cell hyperplasia on infection was also unaffected by conditional KO of ST2 (**Figure 2E-F**). Therefore, susceptibility to *Hpb* infection, and the resulting type 2 immune response to the parasite appeared unchanged in cKO mice.

**Figure 2:**
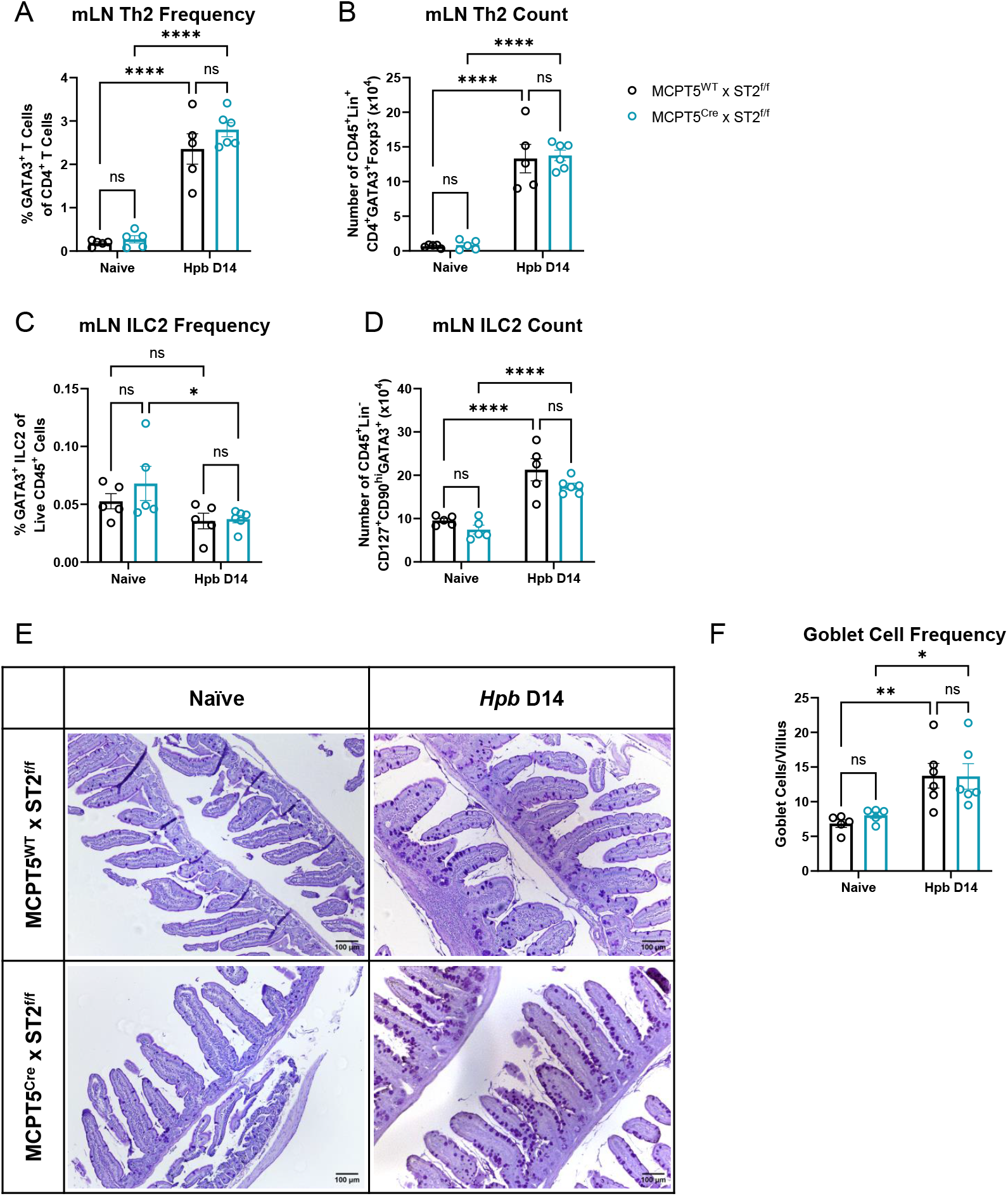
Mast cell ST2 plays no role in type 2 immune responses to *Hpb* infection. MCPT5^C^re x ST2^f/f^ mice and MCPT5^WT^ x ST2^f/f^ mice were infected with 200 L3 *Hpb* larvae by oral gavage, and culled at day 14 of infection, comparing to uninfected naïve mice. Mesenteric lymph node (mLN) cells were taken and stained for flow cytometry to assess proportions (**A**) and numbers (**B**) of Th2 cells, and proportions (**C**) and numbers (**D**) of type 2 innate lymphoid cells (ILC2). Small intestinal Swiss rolls were stained with Periodic acid-Schiff (PAS): representative images shown in (**E**), and enumeration of goblet cells per villus in (**F**). Goblet cells were quantified across 3 ROIs per sample, with at least 6 villi quantified per treatment group. Scale bar represents 100 μm. Data in (**A-D**) is representative of at least 2 experiments, each with n=4-5/group, and was analysed by 2- way ANOVA and Fisher’s LSD test. Data in (**E-F**) is from a single experiment, with n=4- 5/group, and was analysed by 2-way ANOVA and Fisher’s LSD test. Mice were 8-20 weeks old, with both males and females used. Error bars show mean ± SEM.

### Transcriptomics and proteomics of *Hpb*-infected duodenum

In order to determine if changes could be seen in cKO mice at the site of infection, we then conducted transcriptomic and proteomic analysis in naïve and *Hpb*-infected duodenal tissues. Transcript per million (TPM) values for all samples are shown in **Supplementary Table 1**, and iBAQ values are shown in **Supplementary Table 2**.

Using bulk RNAseq of duodenal samples, we compared the overall transcriptional landscape of naïve and infected mice. Using PCA analysis, mice showed separation by infection status but not by genotype (**Figure 3A**). Likewise, PCA analysis of mass spectrometry proteomic results showed a similar effect (**Figure 3B**).

**Figure 3:**
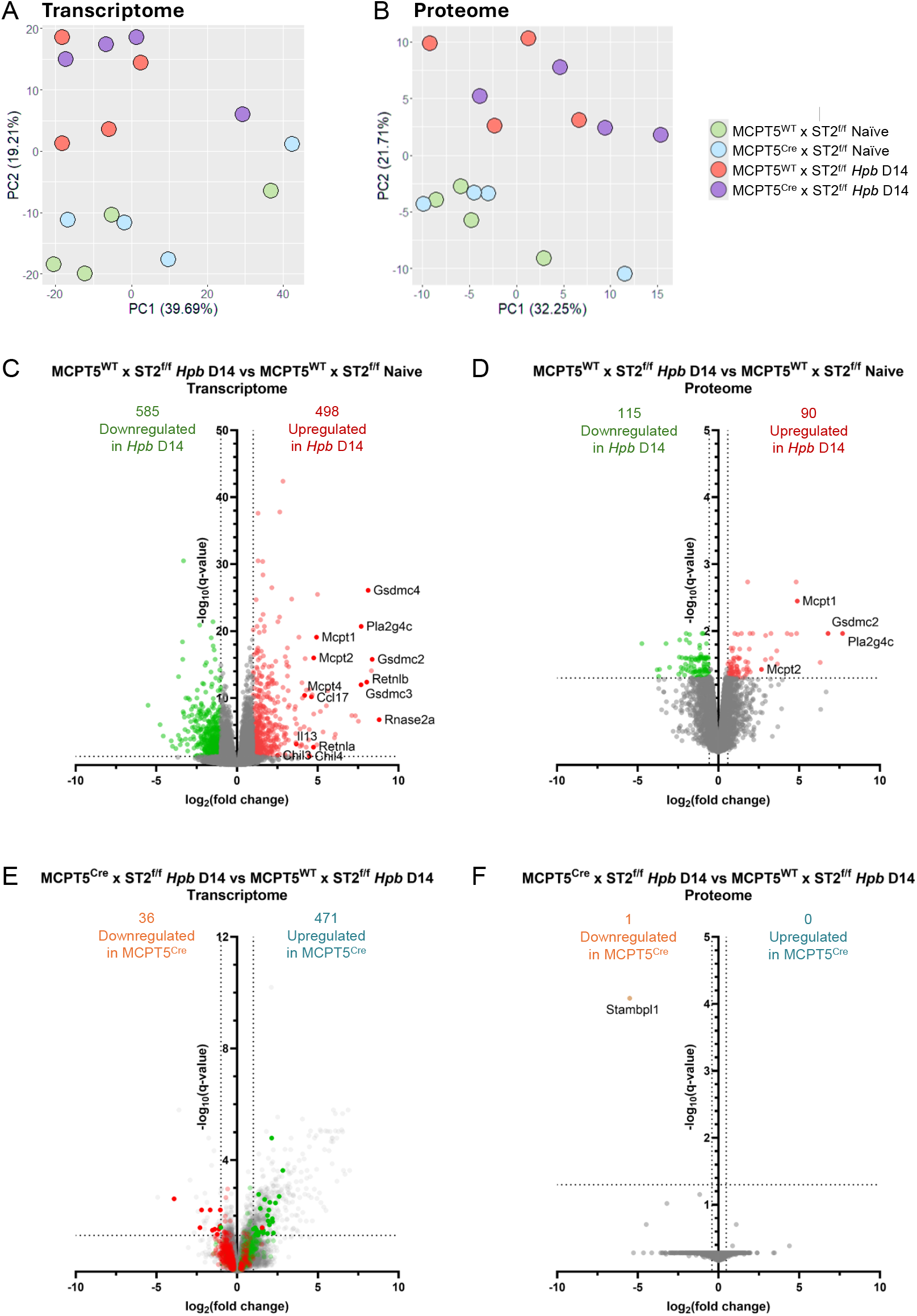
Transcriptomic and proteomic analyses of naïve and *Hpb*-infected MCPT5^C^re x ST2^f/f^ and MCPT5^WT^ x ST2^f/f^ mice. MCPT5^C^re x ST2^f/f^ mice and MCPT5^WT^ x ST2^f/f^ mice were either left naïve or infected with 200 L3 *Hpb* larvae by oral gavage, and culled at day 14 of infection. Duodenal tissue was taken for bulk RNAseq and mass spectrometry analysis. Principal component analysis (PCA) of duodenal bulk RNAseq data (**A**) and mass spectrometry analysis (**B**). Volcano plot of transcript levels in infected versus naïve MCPT5^WT^ x ST2^f/f^ mice, log2(fold change) cutoff = 1, q-value cutoff = 0.05 (**C**). Volcano plot of protein levels in infected versus naïve MCPT5^WT^ x ST2^f/f^ mice, log2(fold change) cutoff = median ± 1 SD, q-value cutoff = 0.05 (**D**). Volcano plot of transcript levels in infected MCPT5^C^re x ST2^f/f^ versus MCPT5^WT^ x ST2^f/f^ mice, log2(fold change) cutoff = 1, q-value cutoff = 0.05 (**E**). Volcano plot of protein levels in infected MCPT5^C^re x ST2^f/f^ versus MCPT5^WT^ x ST2^f/f^ mice, log2(fold change) cutoff = median ± 1 SD, q-value cutoff = 0.05 (**F**). Data is from a single experiment, with n=4-5/group. Mice were 12-20 weeks old, with both males and females used.

**Figure 4:**
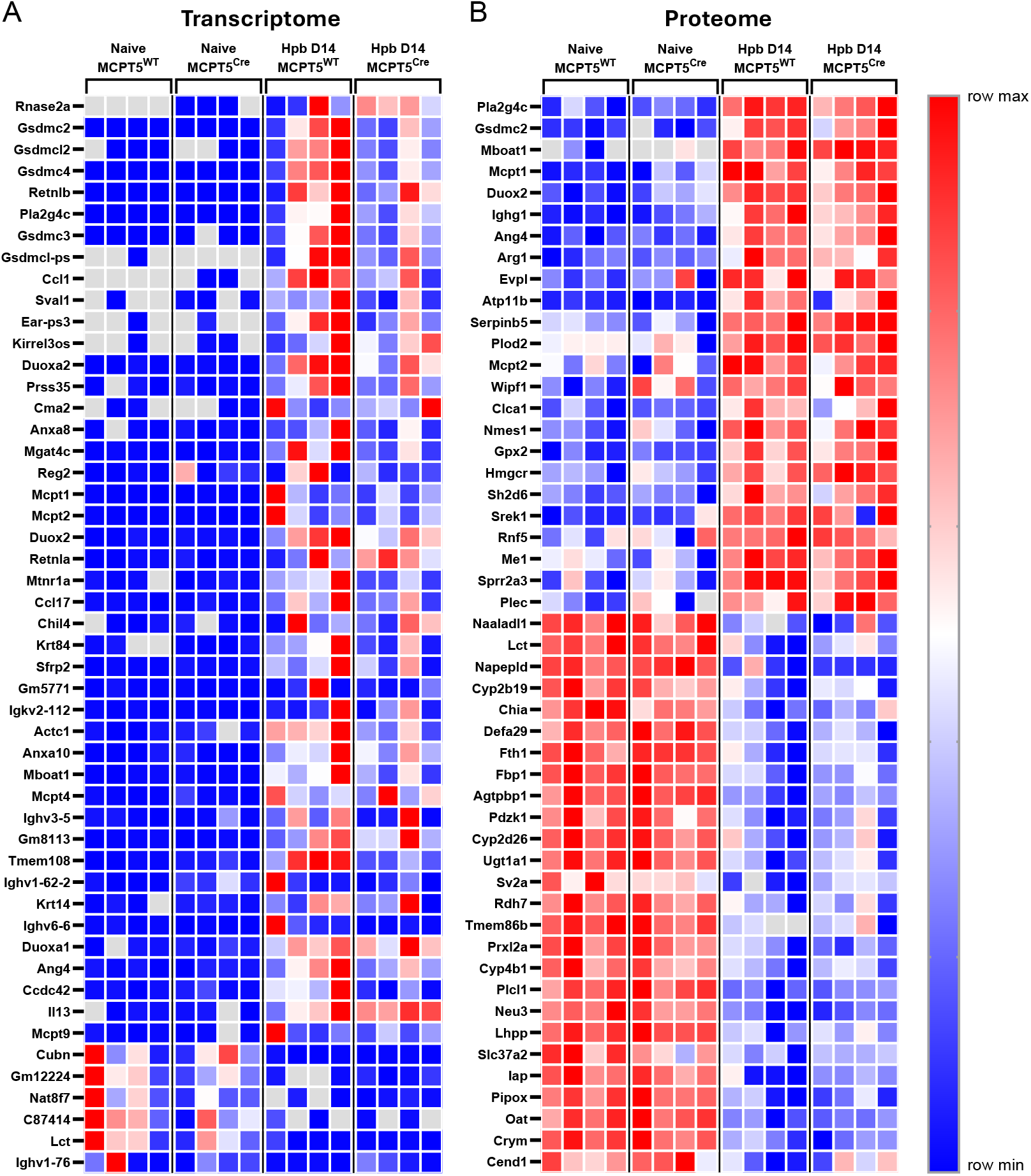
Heatmaps of transcriptomic and proteomic changes in naïve and *Hpb*-infected MCPT5^C^re x ST2^f/f^ and MCPT5^WT^ x ST2^f/f^ mice. Heatmap showing the top 50 significantly differentially expressed genes (A) or proteins (B) by fold change in naïve and *Hpb*-infected mice at day 14 of infection in MCPT5^C^re x ST2^f/f^ mice and MCPT5^WT^ x ST2^f/f^ mice. Grey boxes indicate that gene/protein product was not identified within the sample. Data is from a single experiment, with n=4-5/group. Mice were 12-20 weeks old, with both males and females used.

To first assess the transcriptomic response to *Hpb* infection, bulk RNAseq data of duodenal tissue from naïve control mice and those at day 14 of *Hpb* infection were compared. Infection induced an upregulation of 498 genes and downregulation of 585 genes compared to naïve controls (**Figure 3C**). Strongly upregulated genes included those associated with the type 2 immune response, including mast cell proteases *Mcpt1, Mcpt2* and *Mcpt4*, resistin-like molecules (RELMs) *Retnla* and *Retnlb*, and the cytokines *Il13* and *Ccl17*. In addition, the phospholipase *Pla2g4c* and genes for gasdermin C (*Gsdmc2, Gsdmc3, Gsdmc4*) were also found to be upregulated with *Hpb* infection, similarly to previously published data from other groups (Xi et al., 2021; Yang et al., 2024). Similarly, proteomic analysis replicated many of these effects, showing a significant upregulation of 90 proteins and downregulation of 115 proteins on infection, including upregulation of MCPT1, MCPT2, GSDMC2 and PLA2G4C (**Figure 3D**).

Comparing *Hpb*-infected cKO mice and control mice, cKO mice showed 507 differentially-expressed genes (DEGs): upregulation of 471 genes and downregulation of 36 genes compared to infected littermate controls (**Figure 3E**). However, of these 507 DEGs, only 37 were shared with the 1083 total DEGs in infection. These included 10 genes for which upregulation of expression in infected control mice was blunted in cKO animals (including the genes for galectin-3 and neuregulin-1), while downregulation of 25 genes was likewise blunted in cKOs. When proteomic analysis of cKO versus littermate controls was assessed, only one protein was significantly downregulated in cKO mice (STAMPL1), which was not significantly changed in transcriptomic analysis. Overall, changes in cKO mice were fairly small, and did not show large changes in the type 2 immune response to *Hpb*.

Altogether, these data show large changes in expression in the duodenum during *Hpb* infection, and indicate that ST2 conditional knockout in MCPT5-positive mast cells does not affect susceptibility to *Hpb*, nor the primary type 2 immune response to it.

### Identification of *Hpb* proteins in host tissue

At the day 14 timepoint of *Hpb* infection used here, all parasites have migrated from the wall of the gut into the lumen, where they were flushed out when preparing tissues for analysis after cull. Therefore, we used proteomics to attempt to identify parasite proteins which are present in host tissue at this timepoint, which may represent a subset of parasite secretions that interact with host tissues. To filter out false positives and ensure that only genuine parasite proteins were being detected, we focused downstream analysis on *Hpb* proteins that were: 1) significantly upregulated at day 14 of *Hpb* infection compared to naïve controls; or 2) not identified in any naïve controls but that were present in 3-4 samples from *Hpb*-infected mice. With these criteria, we identified 60 *Hpb* protein hits (**Figure 5, Supplementary Table 3**). Of the 60 hits, 46 were also identified in the excretory/secretory products of *Hpb* (HES). Sequences of the identified proteins were analysed for the presence of a signal peptide, indicating conventional secretion from the parasite (as is the case for the HpARIs, HpBARIs, TGMs and HpBoRB). Of those proteins identified in both the duodenal samples and HES, 18 also contained a signal peptide. These 18 *Hpb* proteins include proteins that are known to be highly expressed and secreted including acetylcholinesterase-1 (AChE-1) and Venom Allergen-Like-1 (VAL-1) (Hewitson et al., 2011). The remaining 14 *Hpb* proteins were identified in *Hpb*-infected duodenal tissue but are not present in HES. These may be proteins that are present within the helminth, identified due to *Hpb* remnants within the duodenum, as evidenced by the presence of a nematode cuticle collagen (Hp_I16296_IG08240_L1042) among them.

**Figure 5:**
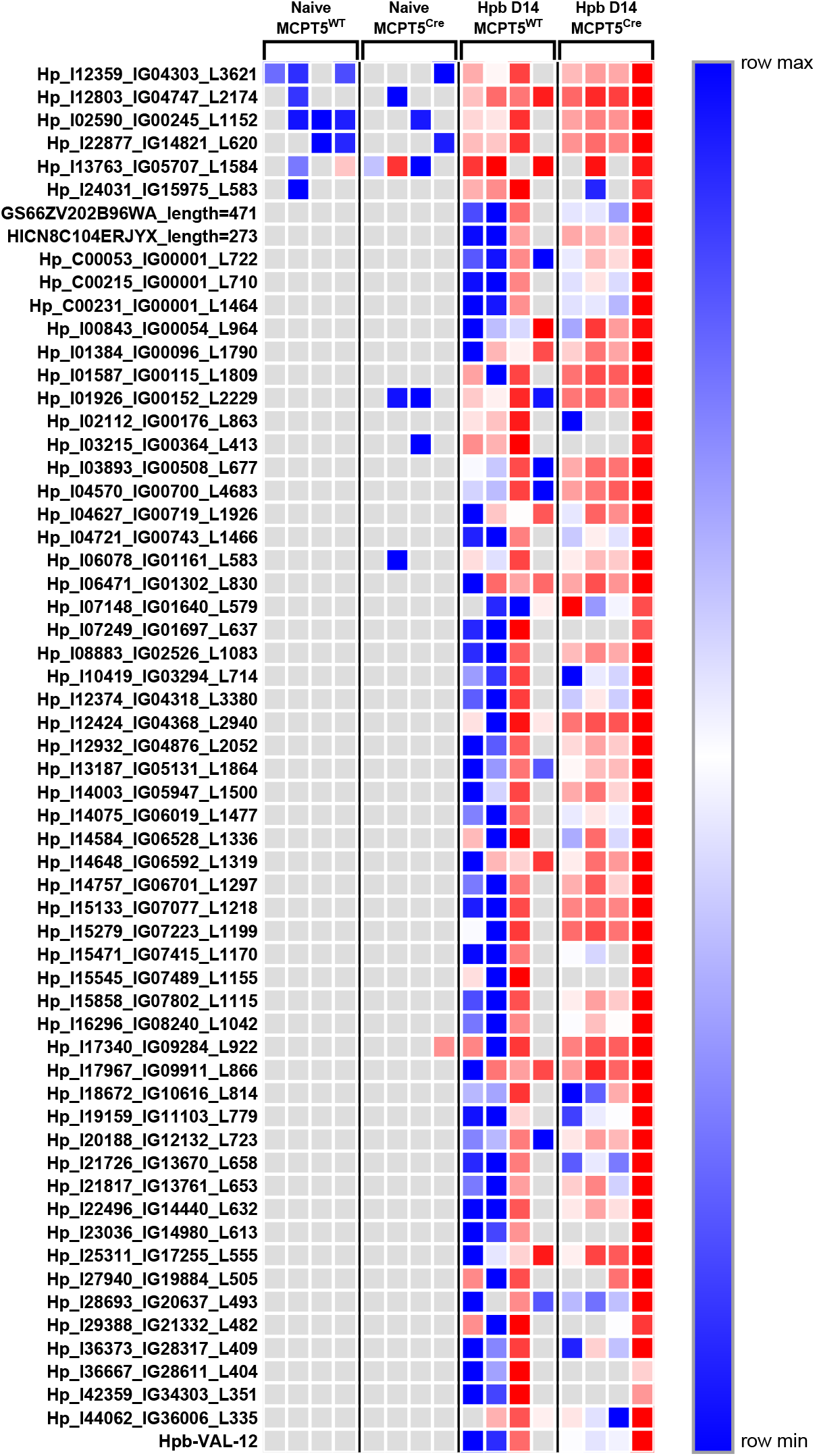
*Hpb* protein products identified in duodenum of infected mice. Heatmap showing abundance of *Hpb* proteins identified in proteome of naïve and *Hpb*- infected mice at day 14 of infection in MCPT5^C^re x ST2^f/f^ mice and MCPT5^WT^ x ST2^f/f^ mice. Grey boxes indicate that protein product was not identified within the sample. Data is from a single experiment, with n=4-5/group. Mice were 12-20 weeks old, with both males and female mice used.

## Discussion

ST2 is known to be an important receptor in the anti-helminth immune response, and the high expression of ST2 on mast cells compared to other immune cells led us to hypothesise a major role for mast cell ST2 in the anti-helminth immune response. In this study we have shown that MCPT5^C^re x ST2^f/f^ mice, which have reduced expression of ST2 on *Mcpt5*-expressing mast cells, show no change in susceptibility to *Hpb* infection compared to littermate controls, suggesting that mast cells may not be the most important cellular responders to IL-33 in helminth infection. Activation of the IL-33/ST2 pathway in other immune cells such as T cells or ILC2s may thus play a more important role in the type 2 immune response in this context. We also use bulk RNAseq and mass spectrometry to assess the host and parasite transcriptomic and proteomic response on infection. We present these datasets as useful resources for the field.

Despite the lack of a detectable role for mast cell-derived ST2, our data does not fully discount the role of mast cells and their expression of ST2 in the immune response against helminths. The mast cell-conditional ST2 knockout mice used in this work lack ST2 expression on mast cells that express *Mcpt5*, which restricts the knockout of ST2 expression to connective tissue mast cells, but not mucosal mast cells. In the intestine, mucosal mast cells express mast cell protease MCPT-1 but not MCPT-5 (Gurish and Austen, 2012), and thus, would retain normal ST2 expression in MCPT5^C^re x ST2f/^f^ mice. Mucosal mast cells, but not connective tissue mast cells, have been found to be important for the timely termination of infection by *Strongyloides ratti*, another intestinal nematode of rodents (Reitz et al., 2017). Mucosal mast cells may thus also play an important role in IL-33-dependent *Hpb* infection, possibly via the IL-33/ST2 pathway.

In this work we also performed transcriptomic and proteomic analysis from duodenal tissues of naïve and *Hpb*-infected cKOs and control mice. Our transcriptomics dataset showed results that corroborate with other published data on intestinal responses during nematode infections (Entwistle et al., 2017; Xi et al., 2021; Campillo Poveda et al., 2025), whereby *Gsdcmc2-4, Pla2g4c* and *Retnlb* were amongst the most upregulated genes at day 14 of *Hpb* infection compared to naïve control mice. In addition, we also provide a matched proteomics dataset that shows upregulation of many of the same gene products.

Mass spectrometry analysis was also used to determine which helminth proteins could be detected within the duodenal tissue of infected mice. This led to us shortlisting 60 *Hpb* proteins of interest which are enriched in infected intestines. Subsequent analysis further narrowed the list down to a subset of 18 proteins that are predicted to contain a signal peptide and have been identified in HES. This subset could contain further immunomodulatory proteins which are interacting with host tissue, and will be the subject of further studies and characterisation. Besides the presence of a signal peptide that suggests a secretory role for the proteins, this subset also includes other proteins known to be abundantly secreted by the parasite, including AChE-1 and VAL-1 (Hewitson et al., 2011). We hypothesise that these proteins may be interacting with the host, and thus staying bound within host tissue even after the parasite has migrated into the lumen. However, known immunomodulatory proteins which bind to host proteins and are present in HES were not found in this dataset, such as HpARI, HpBARI, TGMs and HpBoRB. This could be due to low amount of these proteins secreted by the helminth by day 14 of infection (Pollo et al., 2023), or low local concentration in the duodenal tissue if the proteins diffuse (or are transported) away from the site of deposition, as has been proposed for HpBARI (Smyth et al., 2025).

This study provides evidence against a strong role for connective tissue mast cell responses to IL-33 in helminth infection, and provides a series of datasets looking at parasite and host responses during infection which will be useful for future studies looking at the host-parasite interface and the type 2 immune response to intestinal nematodes.

## Acknowledgements

This work was funded by a Wellcome Investigator award (221914/Z/20/Z) to HJM and a Wellcome PhD Studentship (224014/Z/21/Z) to N.W.P.O. We thank Prof Axel Roers (Heidelberg University Hospital, Germany) for providing *Mcpt5-Cre* mice, and Dr Rebecca Gentek (University of Edinburgh, UK) for advice on their use. Graphic abstract created in BioRender, McSorley, H. (2026) https://BioRender.com/g27lbut. For the purpose of open access, the author has applied a Creative Commons Attribution (CC BY) license to any Author Accepted Manuscript version arising from this submission.

---

**Supplementary Figure 1.**
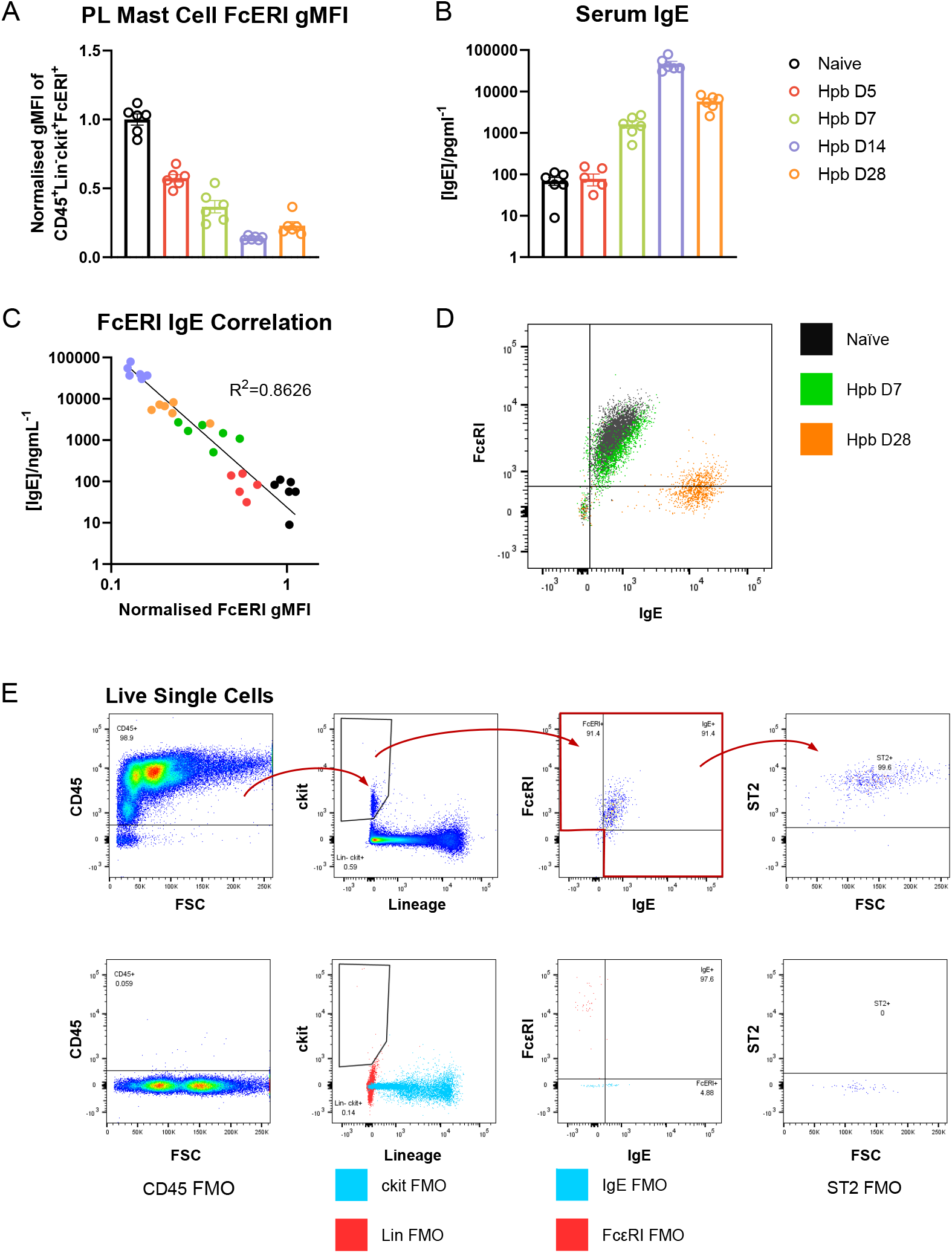
Flow cytometric detection of FcεRI changes across Hpb infection due to IgE-binding. WT C57BL/6J mice were infected with 200 L3 *Hpb* larvae by oral gavage. Separate cohorts of mice were culled at indicated days of *Hpb* infection. Peritoneal lavage (PL) was prepared for flow cytometry to assess mast cell FcεRI expression (geometric mean fluorescence intensity (gMFI) normalised to gMFI in naïve mice) (**A**). Serum levels of total IgE (**B**) were also measured which showed a negative correlation with mast cell FcεRI gMFI (**C**). Data in (**A-C**) is representative of 2 experiments, each with n=6/group. Mice were 8-12 weeks old, with only male WT C57BL/6J mice used. Error bars show mean ± SEM. Flow cytometric analysis of PL mast cells also showed a negative correlation between detection of surface FcεRI and IgE, with representative samples in naïve, *Hpb* D7 and *Hpb* D28 mice shown (**D**). As such, gating strategy of mast cells was updated to include IgE+ cells (**E**).

## Notes

### Competing Interest Statement

The authors have declared no competing interest.

